# Collegiate athlete brain data for white matter mapping and network neuroscience

**DOI:** 10.1101/2020.08.25.267005

**Authors:** Bradley Caron, Ricardo Stuck, Brent McPherson, Daniel Bullock, Lindsey Kitchell, Joshua Faskowitz, Derek Kellar, Hu Cheng, Sharlene Newman, Nicholas Port, Franco Pestilli

## Abstract

We describe a dataset of processed data with associated reproducible preprocessing pipeline collected from two collegiate athlete groups and one non-athlete group. The dataset shares minimally processed diffusion-weighted magnetic resonance imaging (dMRI) data, three models of the diffusion signal in the voxel, full-brain tractograms, segmentation of the major white matter tracts as well as structural connectivity matrices. There is currently a paucity of similar datasets openly shared. Furthermore, major challenges are associated with collecting this type of data. The data and derivatives shared here can be used as a reference to study the effects of long-term exposure to collegiate athletics, such as the effects of repetitive head impacts. We use advanced anatomical and dMRI data processing methods publicly available as reproducible web services at brainlife.io.

## Background & Summary

Elite athletes are highly motivated individuals with physical and psychological characteristics that make them uniquely fit to compete in a sport. Understanding the impact of sports (positive as well as negative) to the brain is an ongoing and active research need^1–26^. Advancing understanding of the effects of sports on the brain is currently hindered by a lack of openly shared datasets and the challenge of collecting data from elite athletes. Our datasets address this challenge by providing magnetic resonance imaging data from 33 athletes and 9 matched controls. As of today, this is the first publicly available and openly shared datasets of college-level athletes.

Relatively recently, efforts to generate large-scale neuroimaging datasets to study the effects of sport on the brain have been funded. Take for example, trackTBI^27^, ENIGMA^28^, and the CARE consortium^29^, which are large scale projects involving multiple investigators and institutions. To contribute to the advancement of science, a good percentage of the data in these projects is or is planned-to-be released for sharing. Because of the administrative burden and funding models, these datasets share data with several restrictions spanning all the way from requirements for co-authorship, study participation or research partnership. Our dataset is different, as we collected a smaller sample of subjects (42) from a single institution and shared the data set openly without administrative overhead. We release this dataset with the intent for it to be used in combination with other datasets, as the size of this dataset alone limits the power for inferences and generalizations.

We share the data utilizing a recently developed unique approach that exploits the free and secure cloud computing platform brainlife.io. The approach integrates, into a single record, both data and reproducible web-services implementing the full processing pipeline^30–32^. The raw data contains both anatomical (T1 weighted) and diffusion-weighted (dMRI) magnetic resonance imaging data. The processed data contains 14 types of derivatives across 42 participants, comprising of 84 brain masks (2 per participant), 42 Freesurfer outputs (1 per participant), 168 fiber orientation distribution (FOD) images (4 per participant), 588 diffusion parameter maps (14 per participant), 42 tractograms (1 per participant), 2,562 segmented major tracts (61 per participant), 756 connectivity matrices (18 per participant), and 42 cortex map data types (1 per participant). The total size of the repository is approximately 676.08 GB of data derivatives comprising 1470 datasets.

The processing pipeline implemented to process this dataset utilizes mainstream neuroimaging software libraries including FSL^33–35^, FreeSurfer^36–55^, MRTrix 3.0^56^, DIPY^57^, and connectome_workbench^58^. The corresponding brainlife.io Apps were developed with a light-weight specification and utilizing modern methods for software containerization making the analyses trackable, reproducible, and reusable on a wide range of computing resources^59^. The present descriptor describes the repository and pipelines published via *brainlife.io* mechanisms. These resources will allow the broader research community to explore the effects of long-term sports participation by exploring high-quality preprocessed neuroimaging, replicate previous examinations of the data, and examine a wide variety of hypotheses without the impediment of the aforementioned barriers.

## Methods

### Data sources

Data is publicly available at https://doi.org/10.25663/brainlife.pub.14.

#### Neuroimaging data sources

Data were collected at the Indiana University Imaging Research Facility (IRF, https://www.indiana.edu/~irf). Data collection was approved by the Indiana University Institutional Review Board.

#### Study participants

A total of fifty-one male participants participated in the study. Twenty-one participants were 4th and 5th-year varsity Indiana University (IU) football “starters” (age 21.1±1.5 years). This number accounts for approximately 60% of the total IU football team active players matching our criteria. Potential participants were excluded if they reported a diagnosed concussion within 6 months of the beginning of the study. The only other exclusion criteria was for safety in the MRI environment. One football player did not complete the study, and three football players did not complete the diffusion MRI (dMRI) scans. This left 17 usable datasets from the Football group. Following scanning, football players received a socioeconomic status survey gathering information regarding estimated family income and the area in which they were raised (i.e. urban, small town, suburbs). Nineteen members of the IU cross-country running team (age 20.2 ± 2.5 years) were included as a non-collision sports group and were matched to the football players based on age and experience level. Three of the players’ anatomical (T1w) images were unusable and thus their data was not included. This left us with 16 eligible members. Our access to the socioeconomic status of non-athlete undergraduates was limited to Psychology and Neuroscience undergraduates who had filled out the socioeconomic survey in an IRB approved subject pool. Eleven controls (non-athletes; age 19.9 ± 3) participants were selected from the limited pool matched on age, sex, estimated family income. None of these individuals were athletes, no additional information on the exercise habits was collected for this group. Two of the controls’ diffusion data contained artifacts that were beyond correction with our processing protocol and thus their data were not included, leaving nine usable datasets. Overall, the released dataset contains usable data from 17 football players, 16 cross-country runners, and 9 non-athletes for analysis (N = 42). In regards to concussion history, two football players had been diagnosed with a concussion approximately 3 years before the study and one player had been diagnosed approximately 2 years before the study. There was no history of concussion in the cross-country runners while at IU. No information was collected regarding the concussion history of the participants before their arrival at IU, however. Although we did not have this information available, we can estimate around 7.25% of football players have been diagnosed with a concussion prior to college given estimates from the literature (Dompier et al., 2015). Participants gave informed written consent that was approved by the Indiana University Institutional Review Board. All participants were recruited through flyers handed out by the athletic trainers of each team or posted around campus. Participants were compensated for participation with a cash payment.

The data detailed here was collected as part of a larger study, which included task-related functional MRI (fMRI) data. The results from the fMRI portion of the study is described in ^60^. Due to this, and the limitations of gathering information via subject pool, focus was placed on collecting neuroimaging data. No other cognitive or behavioral data was collected on the participants, including handedness, IQ, GPAs, or diagnoses of any neuro-cognitive or-developmental disorders.

#### Neuroimaging parameters

Participants were imaged using a 3-Tesla TIM Trio scanner located in the Imaging Research Facility at Indiana University. A 12-channel head coil was used as the 32-channel coil did not fit the heads of our larger subjects. Diffusion-weighted magnetic resonance imaging (dMRI) data were collected with two phase-encoding schemes, i.e anterior-posterior (AP) and posterior-anterior (PA). The following parameters were used for the dMRI pulse sequence: TR/TE = 4930/99.6 ms, iPAT acceleration factor = 2; voxel size = 2×2×2 mm isotropic, 143 diffusion-weighting directions. As student athletes have demanding schedules, emphasis was given to minimizing time of participation when designing the study. Because of this, and the additional fMRI component of the larger study, only two diffusion gradient strengths (b-values) were collected.Sixty-four diffusion gradient directions were collected for each gradient strength,b = 1000 s/mm^2^ and b = 2000 s/mm^2^, respectively. Fifteen non-weighted images were also acquired (b = 0). On T1-weighted (T1w) anatomical image was acquired for each participant using the following sequence: TR/TE = 1800/2.67 ms, TI = 900 ms, flip angle = 9°, bandwidth = 150 Hz/pixel, 160 sagittal slices, FOV = 256 mm, matrix = 256×256, slice thickness = 1 mm, resulting in 1 mm isotropic voxels.

#### Anatomical (T1w) preprocessing

Raw anatomical (T1w) images were preprocessed using the *fsl_anat* functionality provided by the FMRIB Software Library (FSL)^33–35^ implemented as brainlife.io app https://doi.org/10.25663/brainlife.app.273. In brief, the raw anatomical (T1w) images were cropped and reoriented to match the orientation of the MNI152 template. Then, the cropped and reoriented images were *linearly* and *non-linearly* aligned to the MNI152 0.8mm template using *flirt* and *fnirt* respectively^61–63^. The *linearly* aligned images will hereafter be referred to as the ‘acpc aligned’ anatomical (T1w) images. The warps generated from the *non-linear* alignment were subsequently used for mapping diffusion metrics to the cortex^64^. Following alignment, the ‘acpc aligned’ anatomical (T1w) images were processed via Freesurfer’s *recon-all* function to generate pial (i.e. cortical) and white matter surfaces and to parcellate the brain into known anatomical atlases^36–55^ implemented as brainlife.io app https://doi.org/10.25663/bl.app.0. The Destrieux (aparc.a2009s) atlas was used for subsequent white matter tract segmentation and for mapping of diffusion metrics to the cortical surface^65^. For network generation, the multi-modal 180 cortical node parcellation^66^ was mapped to the Freesurfer segmentation of each participant implemented as brainlife.io app https://doi.org/10.25663/bl.app.23. Finally, the ‘acpc aligned’ anatomical (T1w) image was segmented into different tissue-types using MRTrix 3.0’s *5ttgen* functionality^55,56^ implemented as brainlife.io app https://doi.org/10.25663/brainlife.app.239. The gray- and white-matter interface mask was subsequently used as a seed mask for white matter tractography.

Preprocessed anatomical images, and their derivatives, were visually QA’d for common artifacts by BC, RS, and FP. Specifically, the ‘acpc aligned’ anatomical (T1w) images were examined for proper alignment and tissue-contrast. Freesurfer surfaces and parcellations were examined for common surface artifacts and improper voxel identification in parcellations. Any identified issues were manually corrected in Freesurfer and reuploaded before further analysis. The gray- and white-matter interface mask was visually examined for proper separation of the gray- and white-matter in the ‘acpc aligned’ anatomical (T1w) issue.

#### Diffusion (dMRI) preprocessing

Raw dMRI images were first reoriented to match the orientation of the MNI152 template using the fslreorient2std command provided by FSL. The gradients orientation were then checked using MRTrix 3.0’s *dwigradcheck* functionality^67^. Following gradient checking, PCA denoising was performed using MRTrix 3.0’s *dwidenoise* functionality^68^. This was followed by Gibbs deringing using MRTrix 3.0’s *mrdegibbs* functionality^69^. The opposite-facing distortions corresponding to each phase encoding direction (i.e. PA and AP) were then combined into a single corrected image in a method similar to the one described in Andersson and colleagues (2003)^34,70^ (i.e. top-up command) as provided by FSL^33,35^. Eddy-current and motion correction was then applied via the *eddy_cuda8.0* with replacement of outlier slices (*i.e. repol*) command provided by FSL^71–74^. Following this, dMRI images were debiased using ANT’s *n4* functionality^75^ and the background noise not associated with the diffusion signal was cleaned using MRTrix 3.0’s *dwidenoise* functionality^68^. Finally, the preprocessed dMRI images were registered to the ‘acpc aligned’ anatomical (T1w) image using FSL’s *epi_reg* functionality^61–63^ and resliced to 1mm isotropic voxels. The preceding steps were implemented as brainlife.io app https://doi.org/10.25663/bl.app.68. In sum, the dMRI data was interpolated 4 times: 1) following top-up, 2) following eddy, 3) following epi_reg, and 4) during reslicing. These steps were implemented as https://doi.org/10.25663/bl.app.68 Brainmasks of the preprocessed, acpc-aligned dMRI images were then used for subsequent modelling and tractography using FSL’s *bet2* functionality^76^ implemented as brainlife.io App https://doi.org/10.25663/brainlife.app.163.

Quality control was estimated by calculating the Signal to Noise Ratio (SNR) of the diffusing data. To quantify the SNR in the preprocessed, acpc-aligned dMRI data, the workflow provided by Dipy to map SNR in the corpus callosum was used^57,77,78^ implemented as brainlife.io app https://doi.org/10.25663/bl.app.120. SNR values reported are generated from this step.

#### White matter microstructure modeling (DTI)

In order to investigate advanced microstructural properties of white matter, the diffusion tensor (DTI) model was fit to the preprocessed, acpc-aligned dMRI data using FSL’s *dtifit* functionality implemented as brainlife.io app https://doi.org/10.25663/brainlife.app.292. For white matter tract profiles, the default parameters of *dtifit* were used and the b = 1000 shell was chosen for fitting. However, for mapping of the DTI measures to the cortex, both the b = 1000 and b = 2000 shells were used, kurtosis was calculated, and the sum of squared errors was outputted following the parameters used in Fukutomi et al^64^.

### White matter microstructure modeling (NODDI)

In order to investigate advanced microstructural properties of white matter, the Neurite Orientation Dispersion and Density Imaging (NODDI)^79^ model was fit to the multi-shell (i.e. b = 1000, 2000 s/mm^2^) acpc-aligned dMRI data via the Accelerated Microstructure Imaging via Convex Optimization (AMICO; https://github.com/daducci/AMICO; ^80^) toolbox implemented as brainlife.io app https://doi.org/10.25663/brainlife.app.365. The AMICO toolbox was used in order to significantly speed-up the time necessary to fit the NODDI model by reformulating the NODDI model as a linear system, without sacrificing accuracy^80^. For major white matter tract analysis, the isotropic diffusivity parameter (d_iso_) was set to 3.0×10-3 m^2^/s (the rate of unhindered diffusion of water) while the intrinsic free diffusivity parameter (d_//_) was set to 1.7×10-3 mm^2^/s. For cortical white matter parcel analyses, the isotropic diffusivity parameter was also set to 3.0×10-3 mm^2^/s while the intrinsic free diffusivity parameter was set to 1.1×10-3 mm^2^/s, which is the optimal value of diffusivity found by Fukutomi et al^64^.

#### White matter microstructure modeling (CSD)

The CSD model was fit to the preprocessed multi-shell data utilizing MRTrix 3.0 dwi2fod function across 4 maximum spherical harmonic orders (i.e. *L*_max_) parameters (2,4,6,8) implemented as brainlife.io app https://doi.org/10.25663/brainlife.app.238^67,81–83^. *L*_max_’s 6 and 8 were chosen for subsequent white matter tractography, however, all *L*_max_’s are included in the released data.

#### White matter microstructure modeling (Tractography)

Anatomically-constrained probabilistic tractography (ACT)^84^ via MRTrix3’s *tckgen* functionality implemented as brainlife.io app https://doi.org/10.25663/brainlife.app.297 was used to generate tractograms on preprocessed multi-shell dMRI data for each participant. A total of 1.5 million was tracked over both lmax6 and lmax8. The two tractograms were then combined to create a single tractogram of 3 million streamlines via Vistasoft functionality implemented as brainlife.io app https://doi.org/10.25663/brainlife.app.305. The step-size was set to 0.2mm for both lmax6 and lmax8. The minimum length of streamlines was set to 25mm, and the maximum length was set to 250mm. A maximum angle of curvature of 35° was set. The merged tractogram of 3 million streamlines was then used for subsequent white matter tract segmentation and network generation.

#### White matter microstructure modeling (Segmentation & Cleaning)

61 human white matter tracts were segmented using a custom version of the white matter query language^85^ implemented as brainlife.io app https://doi.org/10.25663/brainlife.app.188. These tracts include the following: L/R arcuate, aslant, corticospinal tract (CST), contralateral anterior frontal cerebellar tracts, contralateral motor cerebellar tracts, inferior fronto-occipital fasciculus (IFOF), inferior longitudinal fasciculus (ILF), middle longitudinal fasciculus-angular gyrus (MDLF-ang) and-superior parietal lobule (MDLF-spl), motor cerebellar tracts, occipital cerebellar tracts, superior longitudinal fasciculus components 10026;2 and 3 (SLF-1 & 2, SLF-3), temporal-parietal connection, thalamic cerebellar tracts, uncinate, vertical occipital fasciculus (VOF), Baum’s and Meyers’ loops, cingulum, frontal thalamic tracts, motor thalamic tracts, parietal arcuate (pArc), parietal thalamic tracts, spinothalamic tracts, and temporal thalamic tracts. The callosal tracts, including anterior frontal, forceps major, forceps minor, middle frontal, and parietal corpus callosum, are also included.

Following tract segmentation, outlier streamlines were removed using mba’s *mbaComputeFibersOutliers* functionality^86^ implemented as brainlife.io app https://doi.org/10.25663/brainlife.app.195. For each tract, the spatial ‘core’ representation of the tract was computed by averaging the streamline coordinates across all streamlines in a tract. Streamlines were removed if their length was 4 standard deviations from the length of the ‘core’ representation and/or were located 4 standard deviations away from the ‘core’ representation of the tract. The cleaned segmentations were then used for all subsequent analyses.

#### White matter microstructure modeling (Tract profiles)

Tract profiles^87^ for each DTI parameter estimate (i.e. AD, FA, MD, RD) and NODDI parameter estimate (i.e. NDI, ODI, ISOVF) were generated by estimating the “core” representation of each tract, resampling and segmenting each streamline into 200 equally-spaced nodes, applying a gaussian weight to each streamline based on the distance away from the “core”, and obtaining the weighted average metric at each node. This was performed using MATLAB code utilizing the Compute_FA_AlongFG command provided by Vistasoft (https://github.com/vistalab/vistasoft) implemented as brainlife.io app https://doi.org/10.25663/brainlife.app.361.

#### White matter network modeling (Network generation)

Structural networks were generated using the multi-modal 180 cortical node atlas and the merged tractograms for each participant using MRTrix3’s *tck2connectome*^*88*^ and *tcksift2*^*89*^ functionality implemented as brainlife.io app https://doi.org/10.25663/brainlife.app.394. SIFT2 was used to generate a cross-sectional area weight value for each streamline in order to accurately reflect density. Connectomes were then generated by computing the number of streamlines intersecting each ROI pairing in the 180 cortical node parcellation. Multiple adjacency matrices were generated, including: count, density (i.e. count divided by the node volume of the ROI pairs), length, length density (i.e. length divided by the volume of the ROI pairs), and average and average density AD, FA, MD, RD, NDI, ODI, and ISOVF. Density matrices were generated using the *-invnodevol* option^90^. For non-count measures (length, AD, FA, MD, RD, NDI, ODI, ISOVF), the average measure across all streamlines connecting and ROI pair was computed using MRTrix3’s *tck2scale* functionality using the *-precise* option^91^ and the *-scale_file* option in *tck2connectome*. These matrices can be thought of as the “average measure” adjacency matrices. Before figure generation, nodes in which less than 50% of the participants had a connection were removed.

#### Cortical white matter microstructure modeling (Cortex mapping)

DTI and NODDI measures were mapped to each participant’s cortical white matter parcels following methods found in Fukutomi and colleagues ^64^ using functions provided by Connectome Workbench^58^ implemented as brainlife.io app https://doi.org/10.25663/brainlife.app.379. First, mid-thickness surfaces between the cortical pial surface and white matter surface provided by Freesurfer segmentation were computed using the wb_command -surface-cortex-layer function provided by Workbench command. A Gaussian smoothing kernel (FWHM = ~4mm, σ = 5/3mm) was applied along the axis normal to the surface, and DTI and NODDI measures were mapped using the wb_command -volume-to-surface-mapping function. Freesurfer was used to map the average NODDI parameter estimates to subcortical white matter parcels.

#### Demographics, brain size, body size

We performed multiple one-way ANOVAS between theour groups utilizing the python repository statsmodels’ ols function to test for differences in the following: age, body weight, SNR, average gray-matter cortical thickness, total brain volume, gray-matter cortical volume, and white matter volume. Bonferroni multiple comparisons correction was performed, and all reported p-values were significantly below a corrected p < 0.0083 (0.05 / 6 measures).

### Data visualization

A majority of the images generated for this descriptor were generated using a number of brainlife.io applications utilizing functionality from FSL and DIPY. A list of these Apps include: Generate images of NODDI/DTI, Generate figures of whole-brain tractogram (TCK), Generate images of mask overlaid on DWI, Generate an image of ODF, Generate images of DWI overlaid on T1, Generate images of tissue type masks, Generate images of T1/DWI, and Plot response function. The other images were generated using brainlife.io’s visualization functionality.

## Data Records

The data outputs on brainlife.io are organized using https://brainlife.io/datatypes. These DataTypes allow applications to exchange and archive data. Data outputs can be conveniently downloaded from brainlife.io using the BIDS standard^92^. The data outputs described below can be downloaded at https://doi.org/10.25663/brainlife.pub.14 ^93^. The standard does not yet provide a specification for processed dMRI, tractograms, white matter tracts, and connectivity matrices. The brainlife.io platform will be updated as soon as the BIDS standard fully describes a specification for the models and tractography, tractometry, and network data. For the time being the specification follows the work previous work^30^. We also provide two additional online tables reporting input and output specimens as requested by the Scientific Data guidelines (see Online-only Table 1 and **Supplemental Table 1**).

**Table 1.**
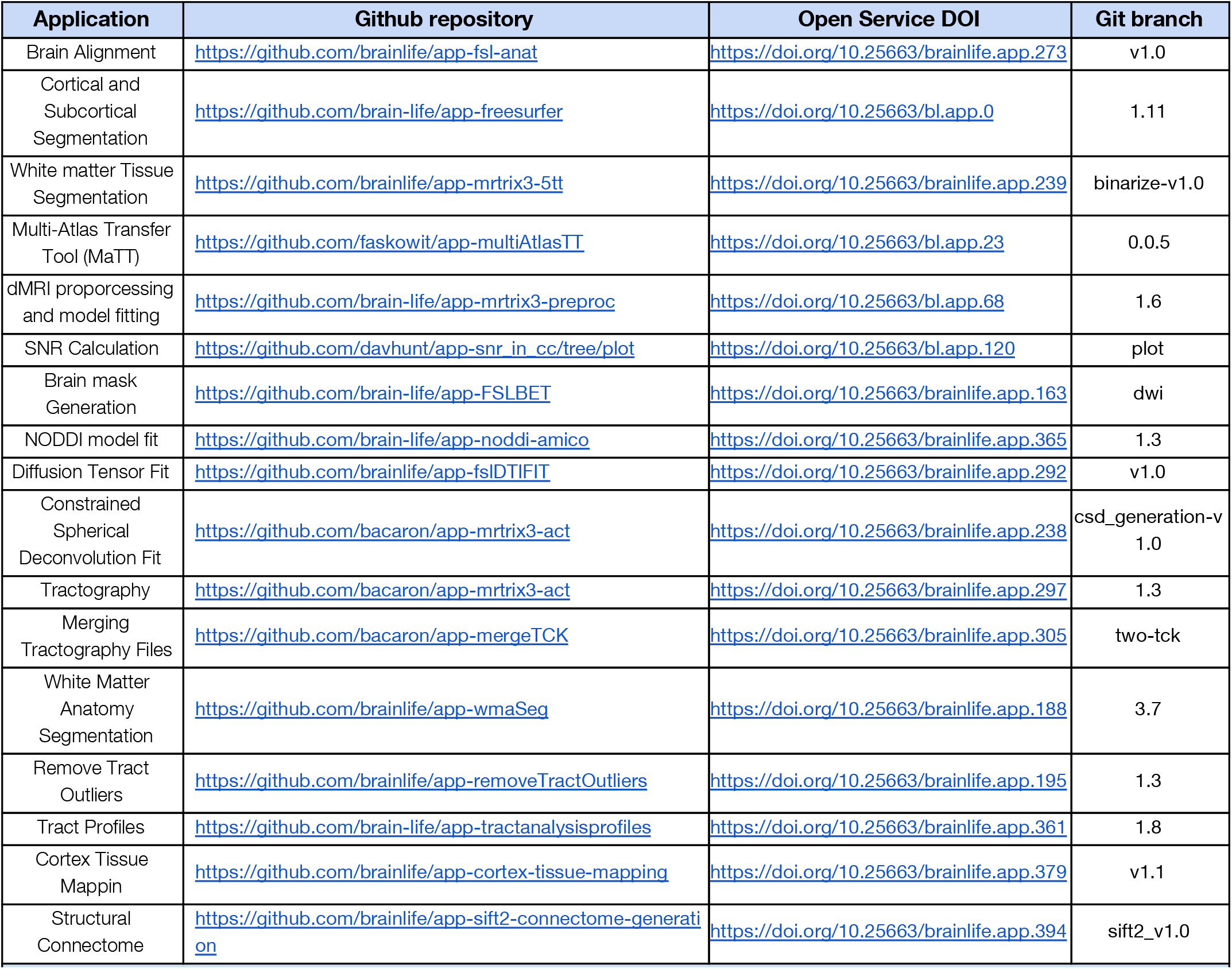
Description and web-links to the open-source code and open cloud services used in the creation of this dataset.

| Application | Github repository | Open Service DOI | Git branch |
| --- | --- | --- | --- |
| Brain Alignment | https://github.com/brainlife/app-fsl-anat | https://doi.org/10.25663/brainlife.app.273 | v1.0 |
| Cortical and Subcortical Segmentation | https://github.com/brain-life/app-freesurfer | https://doi.org/10.25663/bl.app.0 | 1.11 |
| White matter Tissue Segmentation | https://github.com/brainlife/app-mrtrix3-5tt | https://doi.org/10.25663/brainlife.app.239 | binarize-v1.0 |
| Multi-Atlas Transfer Tool (MaTT) | https://github.com/faskowit/app-multiAtlasTT | https://doi.org/10.25663/bl.app.23 | 0.0.5 |
| dMRI preporcessing and model fitting | https://github.com/brain-life/app-mrtrix3-preproc | https://doi.org/10.25663/bl.app.68 | 1.6 |
| SNR Calculation | https://github.com/davhunt/app-snr_in_cc/tree/plot | https://doi.org/10.25663/bl.app.120 | plot |
| Brain mask Generation | https://github.com/brain-life/app-FSLBET | https://doi.org/10.25663/brainlife.app.163 | dwi |
| NODDI model fit | https://github.com/brain-life/app-noddi-amico | https://doi.org/10.25663/brainlife.app.365 | 1.3 |
| Diffusion Tensor Fit | https://github.com/brainlife/app-fslDTIFIT | https://doi.org/10.25663/brainlife.app.292 | v1.0 |
| Constrained Spherical Deconvolution Fit | https://github.com/bacaron/app-mrtrix3-act | https://doi.org/10.25663/brainlife.app.238 | csd_generation-v1.0 |
| Tractography | https://github.com/bacaron/app-mrtrix3-act | https://doi.org/10.25663/brainlife.app.297 | 1.3 |
| Merging Tractography Files | https://github.com/bacaron/app-mergeTCK | https://doi.org/10.25663/brainlife.app.305 | two-tck |
| White Matter Anatomy Segmentation | https://github.com/brainlife/app-wmaSeg | https://doi.org/10.25663/brainlife.app.188 | 3.7 |
| Remove Tract Outliers | https://github.com/brainlife/app-removeTractOutliers | https://doi.org/10.25663/brainlife.app.195 | 1.3 |
| Tract Profiles | https://github.com/brain-life/app-tractanalysisprofiles | https://doi.org/10.25663/brainlife.app.361 | 1.8 |
| Cortex Tissue Mappin | https://github.com/brainlife/app-cortex-tissue-mapping | https://doi.org/10.25663/brainlife.app.379 | v1.1 |
| Structural Connectome | https://github.com/brainlife/app-sift2-connectome-generati on | https://doi.org/10.25663/brainlife.app.394 | sift2_v1.0 |

### T1-weighted Anatomical

T1w image preprocessed and linearly- and nonlinearly-aligned to the MNI152 0.8 MM template using FMRIB Software Library (FSL)’s *fsl_anat* functionality.

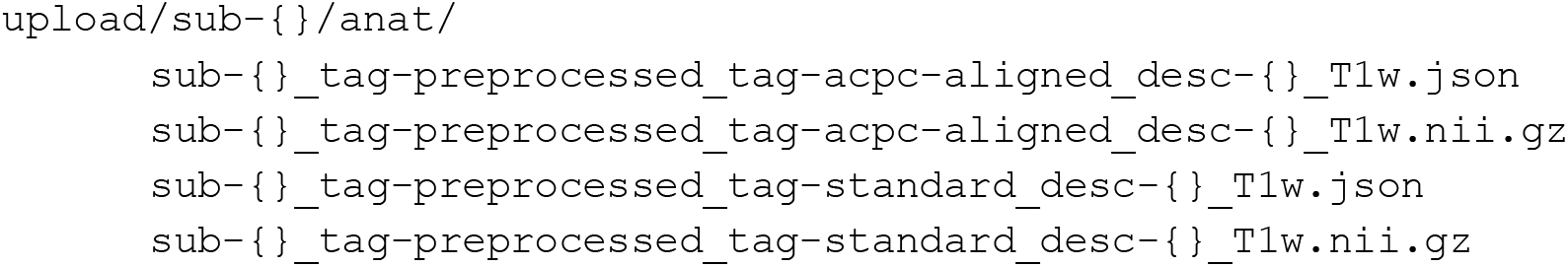

### Non-Linear Image Warping

Warp files describing the non-linear alignment between the raw anatomical (T1w) image and the template image generated from *fsl_anat*. No current BIDS structure exist, https://brainlife.io/ structure is as follows:

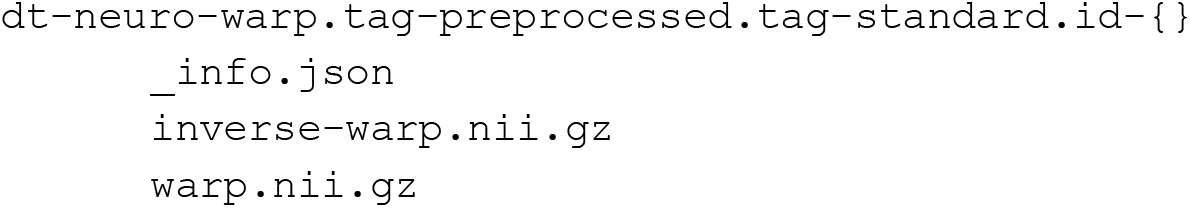

### Freesurfer

Freesurfer output directory containing all derivatives generated during Freesurfer’s *recon-all*. No existing BIDS structure, https://brainlife.io/ structure is as follows:

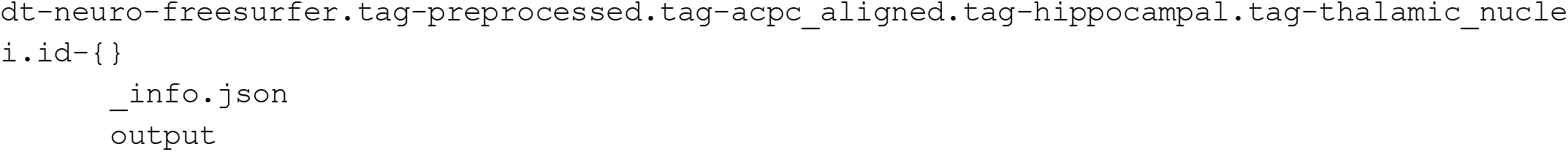

### Multi-Atlas Transfer Tool (MaTT)

The surface and volumated mapping files of the 180 node multimodal parcellation to individual participant surfaces. No existing BIDS structure, brainlife.io structure is as follows:

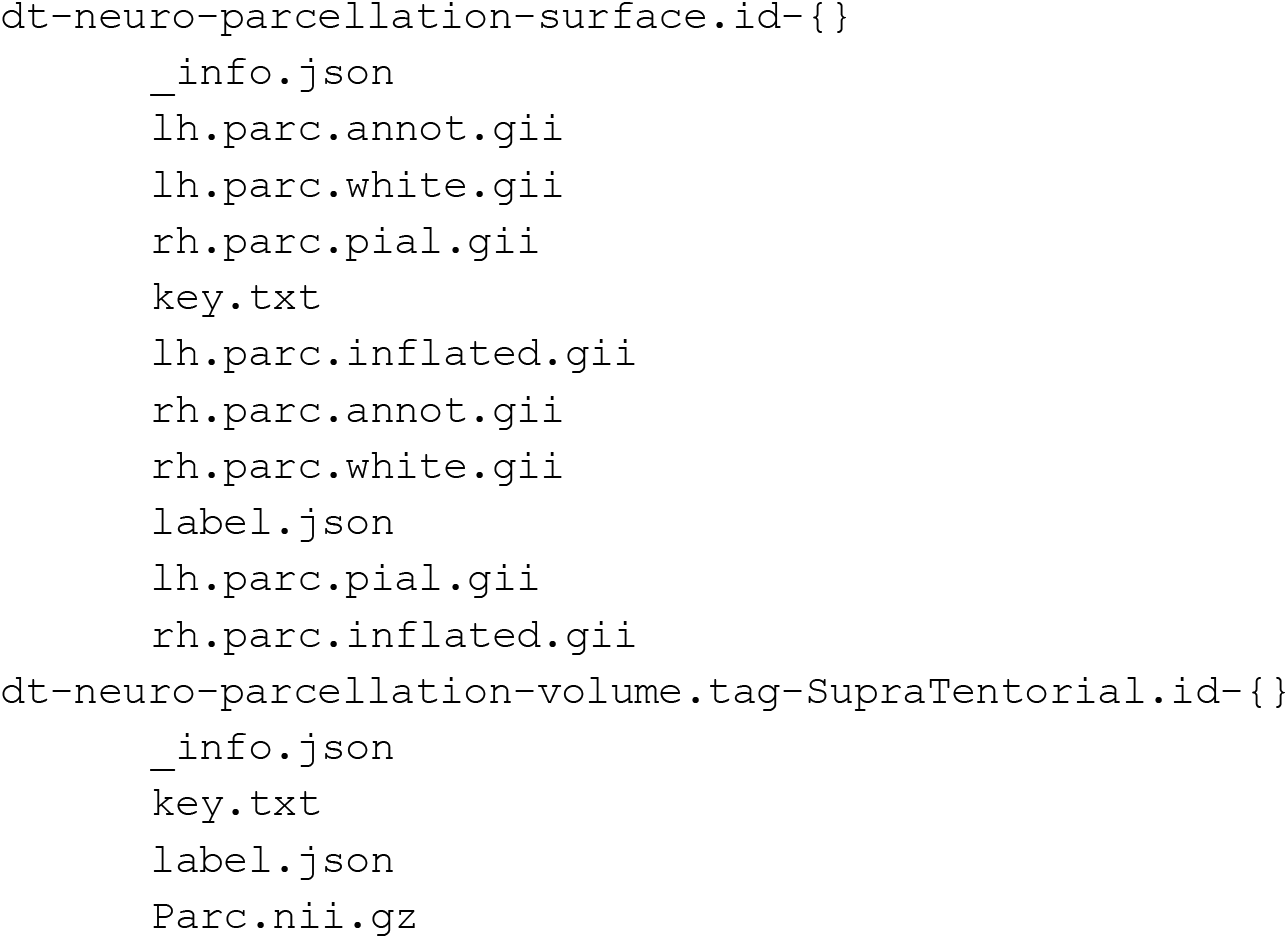

### Tissue-type masks

The 5-tissue type images (GM, WM, CSF, GMWMI) used for tracking. No existing BIDS structure, https://brainlife.io/ structures is as follows:

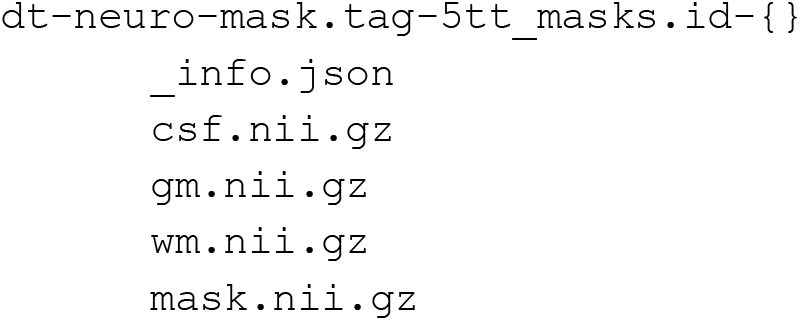

### Diffusion-weighted imaging (dMRI)

The final preprocessed dMRI data used for all further modeling and analyses following FSL Topup & Eddy - CUDA and MRTRix3 preproc.

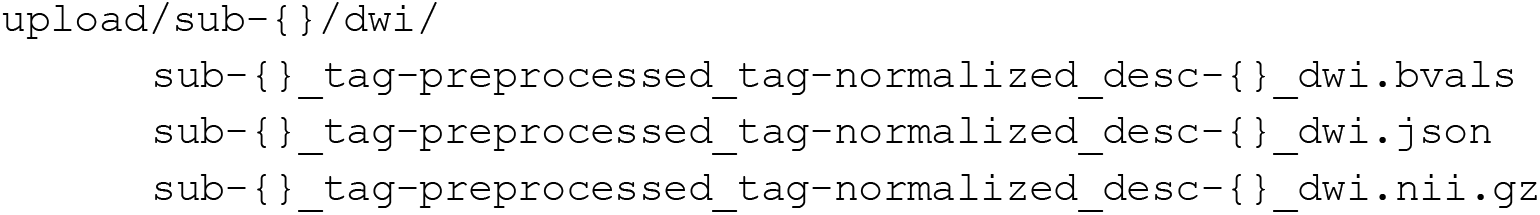

### dMRI Brainmask

The final dMRI brain mask used for all modeling and analyses. No existing BIDS structure, https://brainlife.io/ structures are as follows:

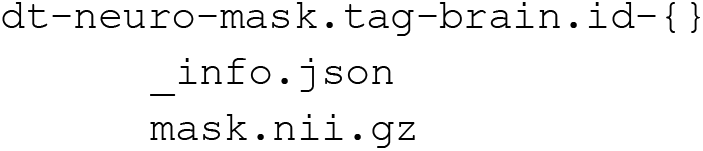

### Neurite Orientation Dispersion Density Imaging (NODDI)

The neurite density, orientation dispersion, and isotropic volume fraction maps generated from NODDI AMICO. No existing BIDS structure, https://brainlife.io/ structure is as follows:

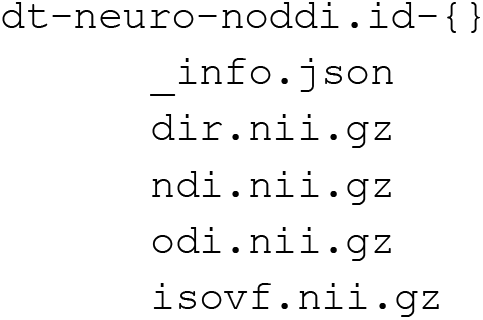

### Diffusion Tensor Imaging (DTI)

The fractional anisotropy, mean diffusivity, axial diffusivity, and radial diffusivity maps generated from DTIFIT.

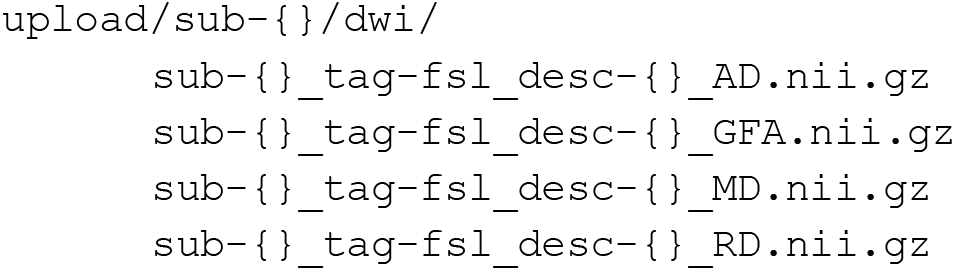

### Constrained Spherical Deconvolution (CSD)

CSD models fit across *L*_max_=2,4,6, and 8 using Fit Constrained Spherical Deconvolution Model For Tracking. No existing BIDS structure, https://brainlife.io/ structure is as follows:

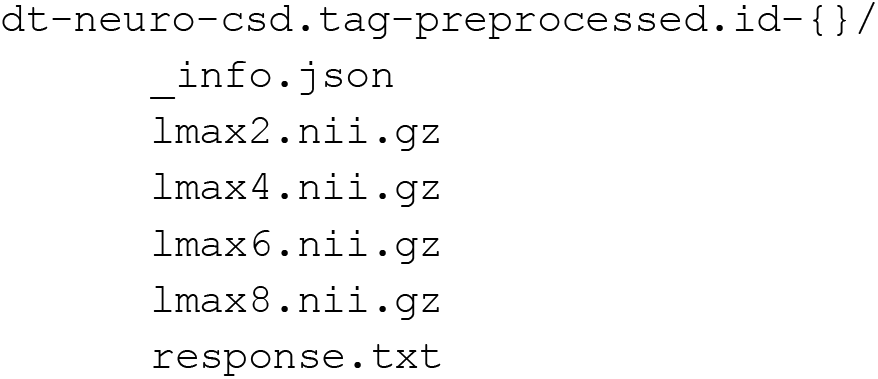

### Tractograms

The merged tractograms across *L*_max_=6 and *L*_max_=8, totaling 3 million streamlines generated from the Anatomically-constrained tractography (ACT) app.

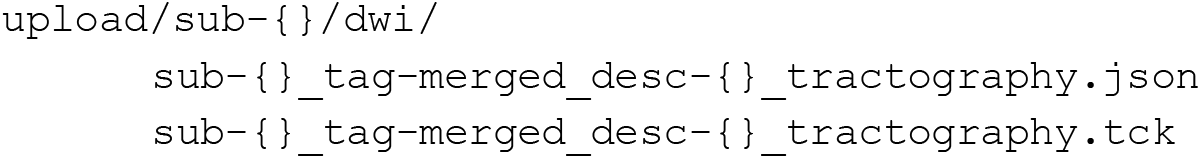

### White Matter Anatomy (wma) Segmentation

Major track segmentation generated from White Matter Anatomy Segmentation. No existing BIDS structure, https://brainlife.io/ structure is as follows:

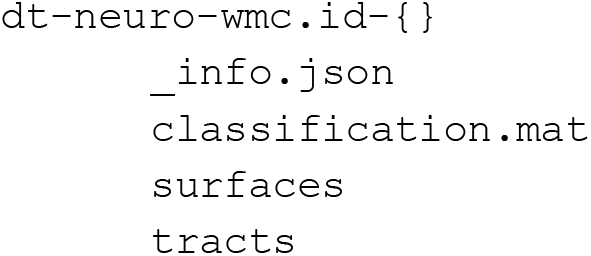

### Segmentation Cleaned

The cleaned tracks from Remove Fiber Outliers. No existing BIDS structure, https://brainlife.io/ structure is as follows:

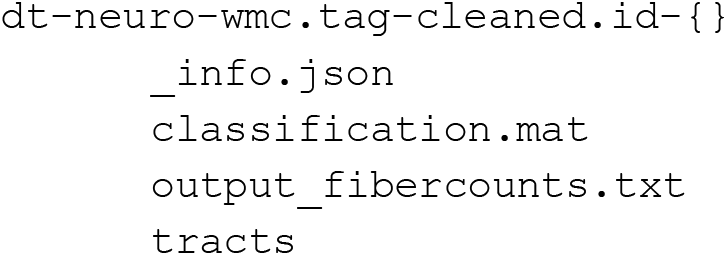

### Tract Profiles

Mapping of DTI and NODDI metrics along the core of the segmented white matter tracts using the Tract Analysis Profiles app. No existing BIDS structure, https://brainlife.io/ structure is as follows:

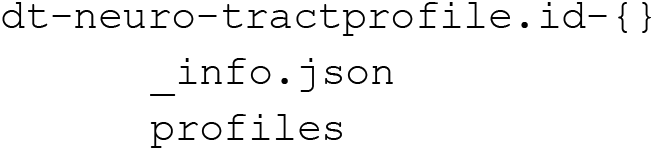

### Network Generation

Network adjacency matrices were generated using the Structural Connectome MRTrix3 (SCMRT) (SIFT2) app. No existing BIDS structure, https://brainlife.io/ structure as follows:

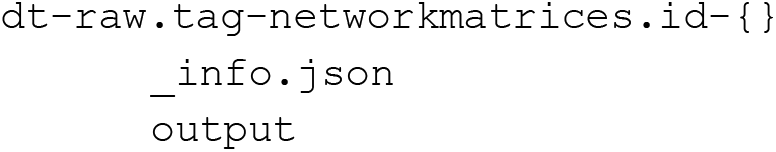

### Cortical DTI and NODDI mapping

Surfaces and DTI and NODDI measure files mapped to the surface generated from Cortical Tissue Mapping. No existing BIDS structure, https://brainlife.io/ structure is as follows:

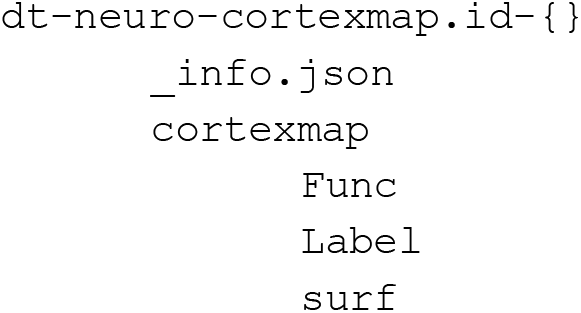

## Technical Validation

In this section, we provide a qualitative evaluation of the data derivatives made available on https://brainlife.io/. We provide qualitative analysis of the preprocessing of the anatomical (T1w) image, including the seed mask, and pial and white matter surfaces generated from Freesurfer. Qualitative images of the dMRI preprocessing, dMRI modeling (CSD and NODDI), dMRI tractography, network generation, and mapping of diffusion measures to the cortical surface are also provided. We further provide a quantitative analysis of the SNR of the dMRI data following preprocessing.

### Anatomical (T1w) preprocessing

Anatomical (T1w) images were *linearly* aligned to the MNI152 0.8 mm template and further segmented into gray-matter, white-matter, CSF, and gray- and white-matter interface masks using https://doi.org/10.25663/brainlife.app.273. See **Methods: Anatomical (T1w) preprocessing** for more details. Each participant’s aligned anatomical images (T1w), and all derivatives generated from the aligned images, are provided.

Figure 1a exemplifies the quality of the linear alignment obtained with https://doi.org/10.25663/brainlife.app.300 in representative participants from each athlete group (i.e. Football: *top*, Cross-Country: *middle*, and Non-Athlete: *bottom*). The gray- and white-matter interface mask (1b) and white matter boundary (1c) are overlaid on the ‘acpc-aligned’ anatomical (T1w) image to further provide quality assurance. These images were generated with https://doi.org/10.25663/brainlife.app.312.

**Figure 1.**
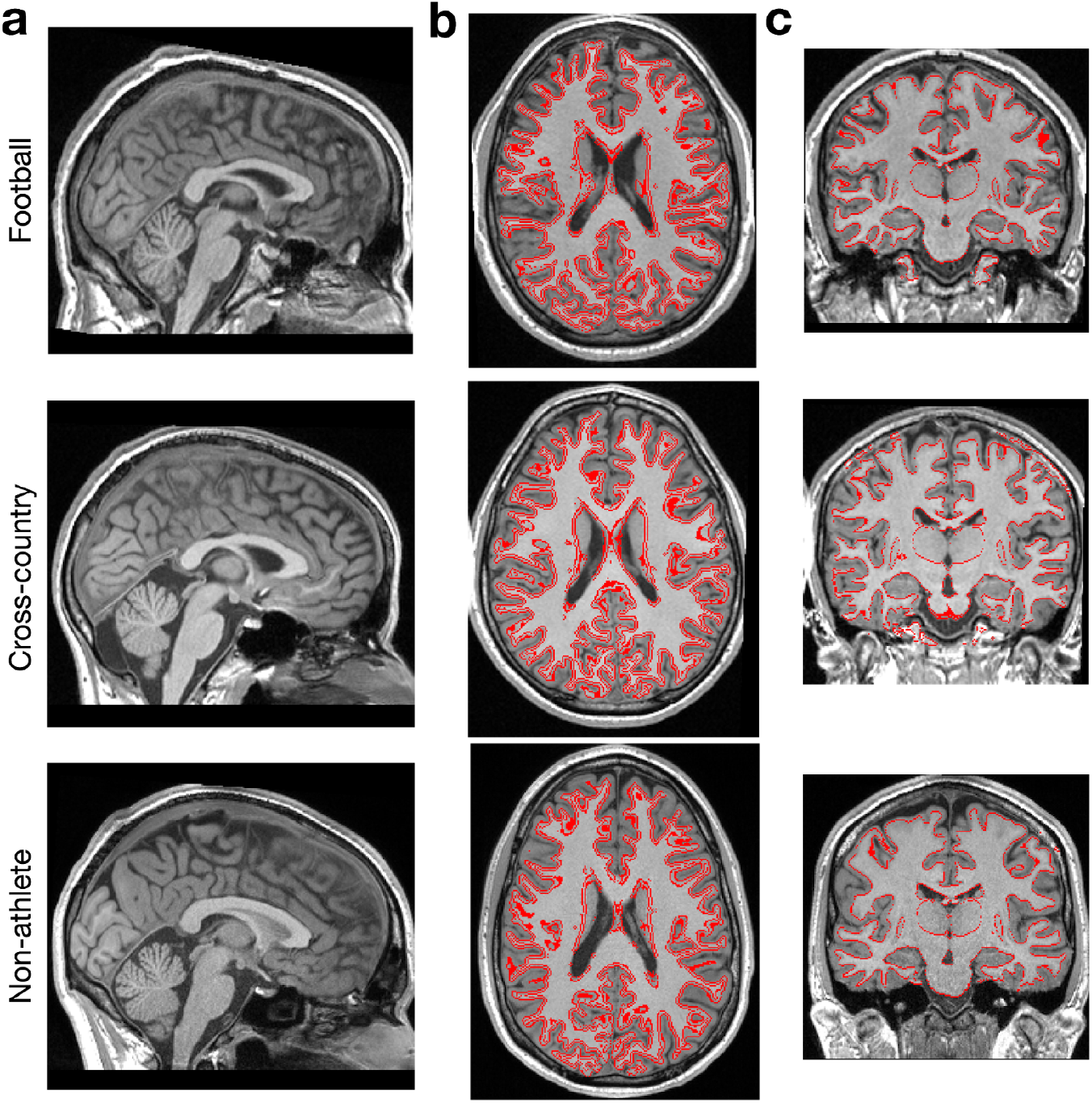
Anatomical (T1w) preprocessing: Alignment and segmentation. Representative mid-plane images from each group of the preprocessed anatomical (T1w) image. (a) Sagittal slice of MNI152 (ACPC) aligned T1 used for all subsequent preprocessing, (b) Axial slice with gray - and white-matter interface outlined (red) used as a seed mask for tractography, and (c) Coronal slice with white-matter boundary outlined (red). Images were generated using https://doi.org/10.25663/brainlife.app.300 and https://doi.org/10.25663/brainlife.app.312

Following alignment and segmentation, Freesurfer was used to generate cortical and white matter surfaces, along with the Destrieux Atlas parcellation using https://doi.org/10.25663/bl.app.0. Figure 2a demonstrates the quality of the surface generation representative participants from each athlete group (i.e. Football: *top*, Cross-Country: *middle*, and Non-Athlete: *bottom*). Images of the Destrieux (aparc.a2009s) atlas^65^ on the pial surface, along with images of the pial and white matter surface outlines overlaid on the ‘acpc-aligned’ anatomical (T1w) image, are provided as a means of quality assurance. Figure 2b illustrates the mapping of the 180 node multimodal atlas^66^ to representative participants from each group mapped using https://doi.org/10.25663/bl.app.23. These images were generated using brainlife.io’s Freeview and Connectome Workbench viewers.

**Figure 2.**
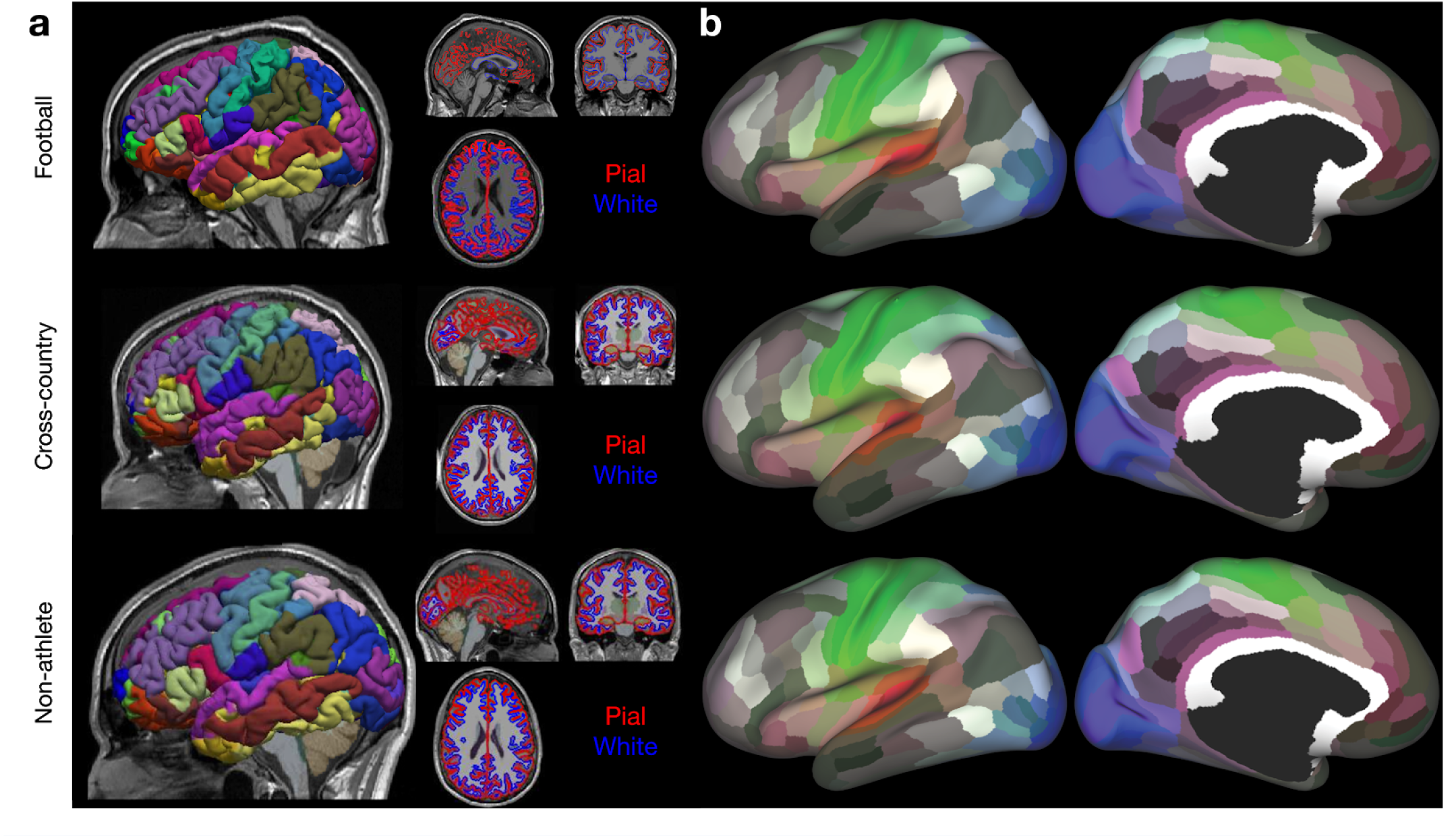
Anatomical (T1w) preprocessing: Freesurfer and 180 node multimodal atlas mapping. **a.** Representative images from each group of the Freesurfer outputs: pial (*red*) and white (*blue*) matter surfaces, and the aparc.a2009s+aseg (i.e. Destrieux) parcellation. Images were generated using brainlife.io’s Freeview viewer. **b.** Representative images from each group of the 180-node multimodal (hcp-mmp) atlas mapped to an inflated representation of the cortical surface. Images were generated using brainlife.io’s Connectome Workbench viewer.

### Diffusion (dMRI) preprocessing

Raw dMRI images were corrected for Gibbs ringing, susceptibility-weighting, eddy currents, motion, biasing, and Rician background noise using a combination of methods. Following preprocessing, the dMRI images were aligned to the ‘acpc-aligned’ anatomical (T1w) image. See **Methods: Diffusion (dMRI) preprocessing** for more details. The preprocessing was performed using https://doi.org/10.25663/bl.app.68. Following preprocessing, the signal-to-noise ratio (SNR) was computed for each subject in the non-diffusion weighted volumes (i.e. b=0) and the diffusion-weighted volumes (i.e. b=1000,2000) separately as a means for quality assurance. The SNR was computed using https://doi.org/10.25663/bl.app.120.

Figure 3a demonstrates the quality of alignment of the dMRI and ‘acpc-aligned’ anatomical (T1w) image from representative participants from each group. The fractional anisotropy (FA) map (see **Methods: White matter microstructure: DTI** for more details on DTI fitting) from each subject is overlaid in red-yellow on the ‘acpc-aligned’ anatomical (T1w) images. Overall, the alignments of the dMRI and the anatomical image are anatomically-sound. The fully preprocessed dMRI images from each participant, along with their corrected b-vectors and b-values, are provided. The images were generated using https://doi.org/10.25663/brainlife.app.309. Figure 3b documents the non-diffusion weighted and diffusion-weighted SNRs for each participant. The average SNR for Football players (*orange*) following preprocessing was 28.354 (± 5.772 SD) for non-diffusion weighted volumes. This was slightly lower than the SNR for Cross-country runners (*pink*) with an average SNR in the non-diffusion weighted volumes of 34.944 (±4.594 SD). Non-athletes overall had the lowest average SNR in the non-diffusion weighted volumes (23.002±7.784 SD).

**Figure 3.**
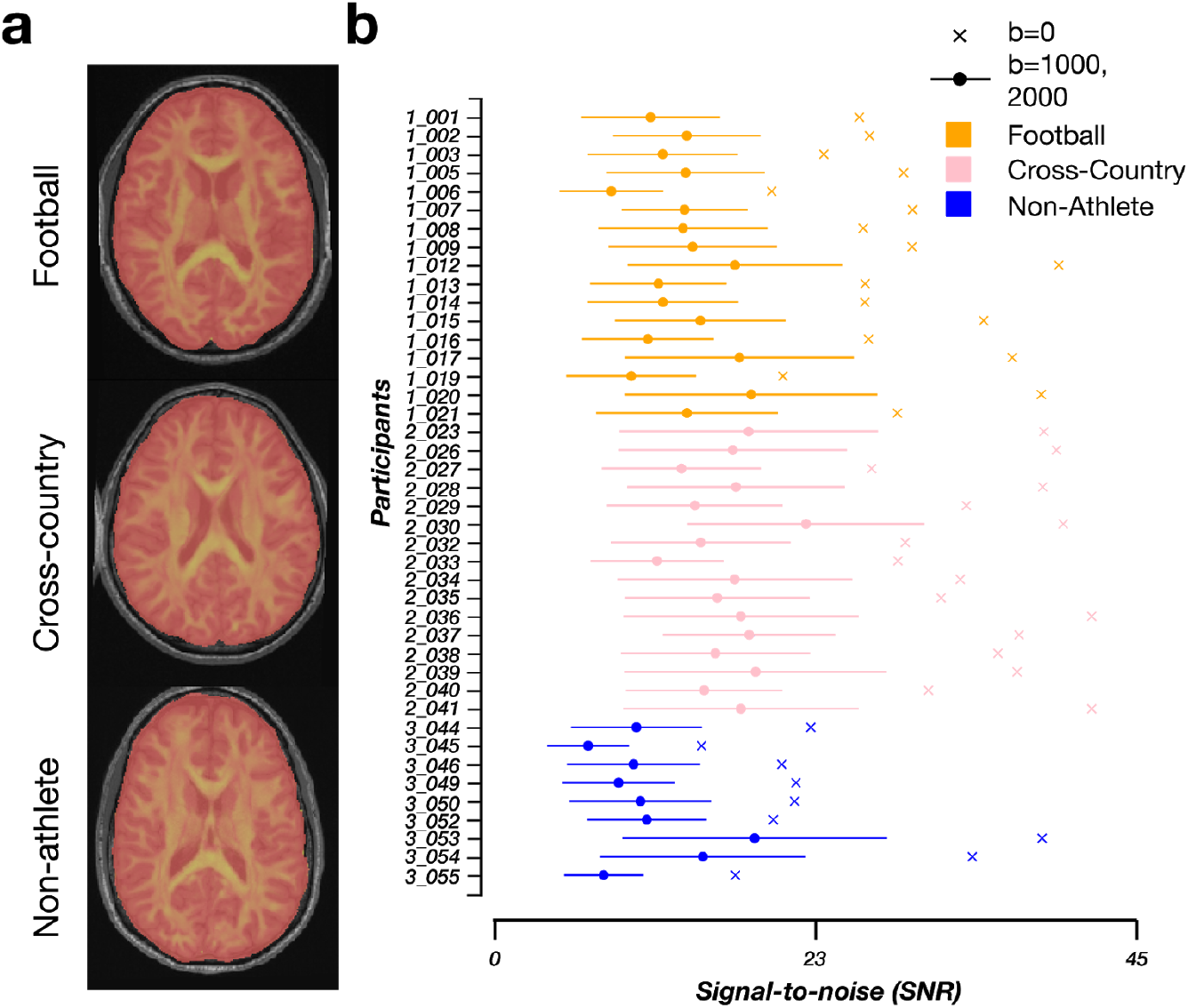
Qualitative and quantitative quality-assurance measures following dMRI preprocessing. **a.** Representative mid-axial images of the fractional anisotropy (FA) (*red-yellow*) image overlaid on the ‘acpc aligned’ anatomical (T1w) image. Images were generated using https://doi.org/10.25663/brainlife.app.309. **b.** The signal-to-noise ratio for each participant from each group (football: *orange*, cross-country: *pink*, non-athlete: *blue*) following dMRI preprocessing. The average SNR was computed in the Corpus Callosum across the b=0 shells (*crosses*) and b=1000,2000 shells (*circles*). Standard deviation across the non-b=0 shells are plotted as error bars for each participant.

### White matter microstructure modeling: DTI and NODDI

Following preprocessing, models of microstructure were fit to the dMRI images. Specifically, the diffusion tensor (DTI) and neurite orientation dispersion density imaging (NODDI) models were fit to the b=1000 and b=1000, 2000 shells respectively. See **Methods: White matter microstructural modeling: DTI & NODDI** for more details. DTI was fit using https://doi.org/10.25663/brainlife.app.292, while NODDI was fit using https://doi.org/10.25663/brainlife.app.365. The DTI and NODDI maps for each participant are provided. Figure 4 demonstrates the quality of fit of both the DTI and NODDI models on representative participants from each group. Specifically, mid-axial slices of the fractional anisotropy (FA),mean diffusivity (MD), axial diffusivity (AD), and radial diffusivity (RD) from the DTI model, and the orientation dispersion and neurite density indices (ODI, NDI) and the isotropic volume fraction (ISOVF) from the NODDI model are presented. There is a high anatomical correspondence between the measures and known anatomical properties. For example, the ventricles across all three participants are saturated in the MD maps, as water moves maximally isotropically. In the white matter, FA and NDI are highest in the highest concentrations of white matter, while ODI is lowest. The images were generated using https://doi.org/10.25663/brainlife.app.302 and https://doi.org/10.25663/brainlife.app.367.

**Figure 4.**
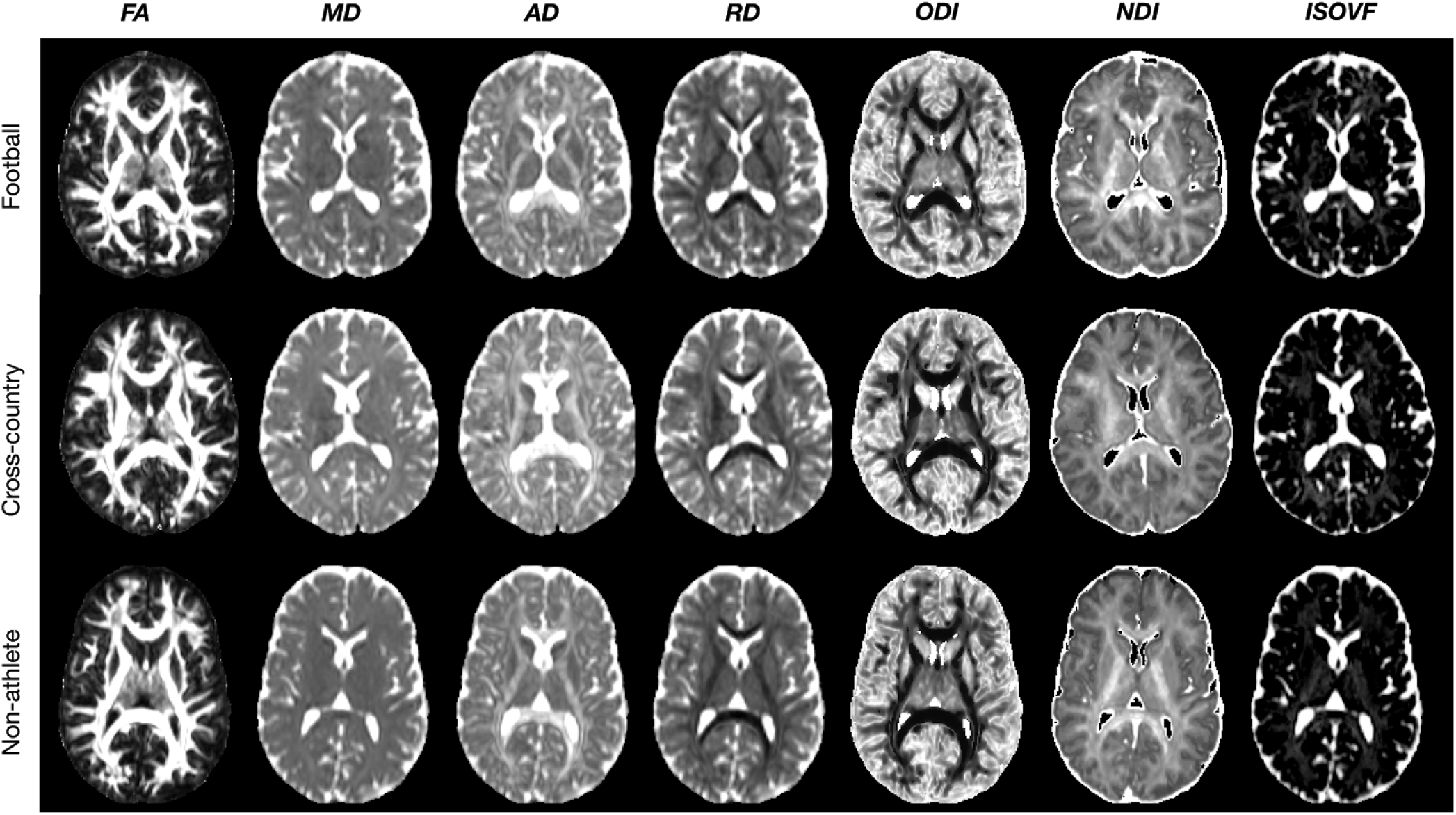
dMRI Modelling: DTI and NODDI. Qualitative quality-assurance figures following fitting of the Diffusion Tensor (DTI) and Neurite Orientation Dispersion Density Imaging (NODDI) models. (a) Representative mid-axial images of the fractional anisotropy (*FA*), mean diffusivity (*MD*), axial diffusivity (*AD*), radial diffusivity (*RD*), orientation dispersion index (*ODI*), neurite density index (*NDI*), and isotropic volume fraction (*ISOVF*). Images were generated with https://doi.org/10.25663/brainlife.app.302 and https://doi.org/10.25663/brainlife.app.367.

### White matter microstructural modeling: CSD

In order to map white matter macrostructure via white matter tractography, the constrained spherical deconvolution model (CSD) was fit to each participant across 4 maximum spherical harmonic orders (i.e. *L*_max_): 2,4,6 and 8. *L*_max_=6,8 were chosen for tracking. See **Methods: White matter microstructural modeling (CSD)** for more details. The CSD was fit using https://doi.org/10.25663/brainlife.app.238. Each participant’s CSD fits across all four *L*_max_’s are provided. Figure 5 demonstrates the quality of fit of the CSD model on representative participants from each group using *L*_max_=8. In the left column, the response function generated is mapped to a sphere, while the right column corresponds to the fiber orientation distribution function (fODF). The response functions demonstrate a quality fit due to the relatively flat shape and sharp folding in the center. In the fODF maps, clear anatomy is distinguished in regions of the highest white matter concentration. Images were generated using https://doi.org/10.25663/brainlife.app.311 and https://doi.org/10.25663/brainlife.app.317.

**Figure 5.**
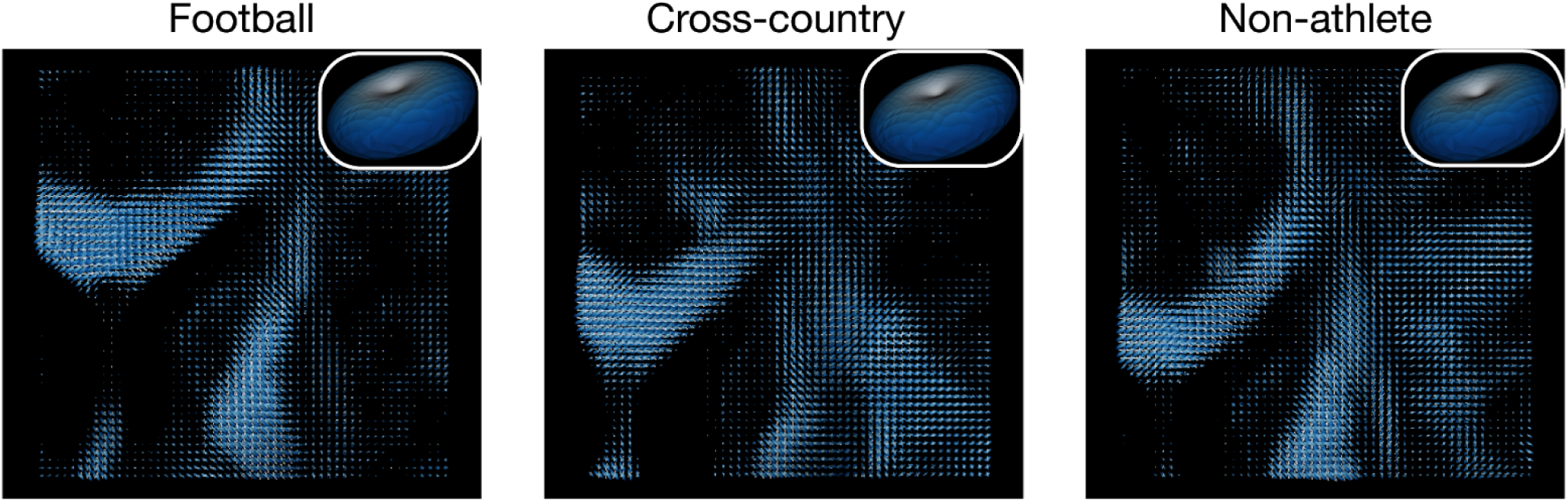
dMRI Modelling: Constrained Spherical Deconvolution. Example outputs of the fiber orientation distribution function (fODF) for maximum spherical harmonic order (*L*_max_) of 8 and the csd response functions (*inset*). Images were generated using https://doi.org/10.25663/brainlife.app.311 and https://doi.org/10.25663/brainlife.app.317.

### White matter microstructure modeling: Anatomically-constrained tractography

Following the fitting of the CSD model to each participant, anatomically-constrained tractography (ACT) was performed on *L*_max_=6,8 to generate whole-brain tractograms containing 3 million streamlines. These tractograms were used for subsequent segmentation and network generation. See **Methods: White matter microstructure modeling (Tractography)** for more details. The tractograms were generated using https://doi.org/10.25663/brainlife.app.297, and merged using https://doi.org/10.25663/brainlife.app.305. Each participant’s whole-brain tractogram containing 3 million streamlines per tractogram is provided. Figure 6a demonstrates the quality of the tractography in representative participants from each group. The whole-brain tractogram of each participant shows high densities of streamlines filling the entire brain volume, as expected. The images were generated using https://doi.org/10.25663/brainlife.app.310.

**Figure 6.**
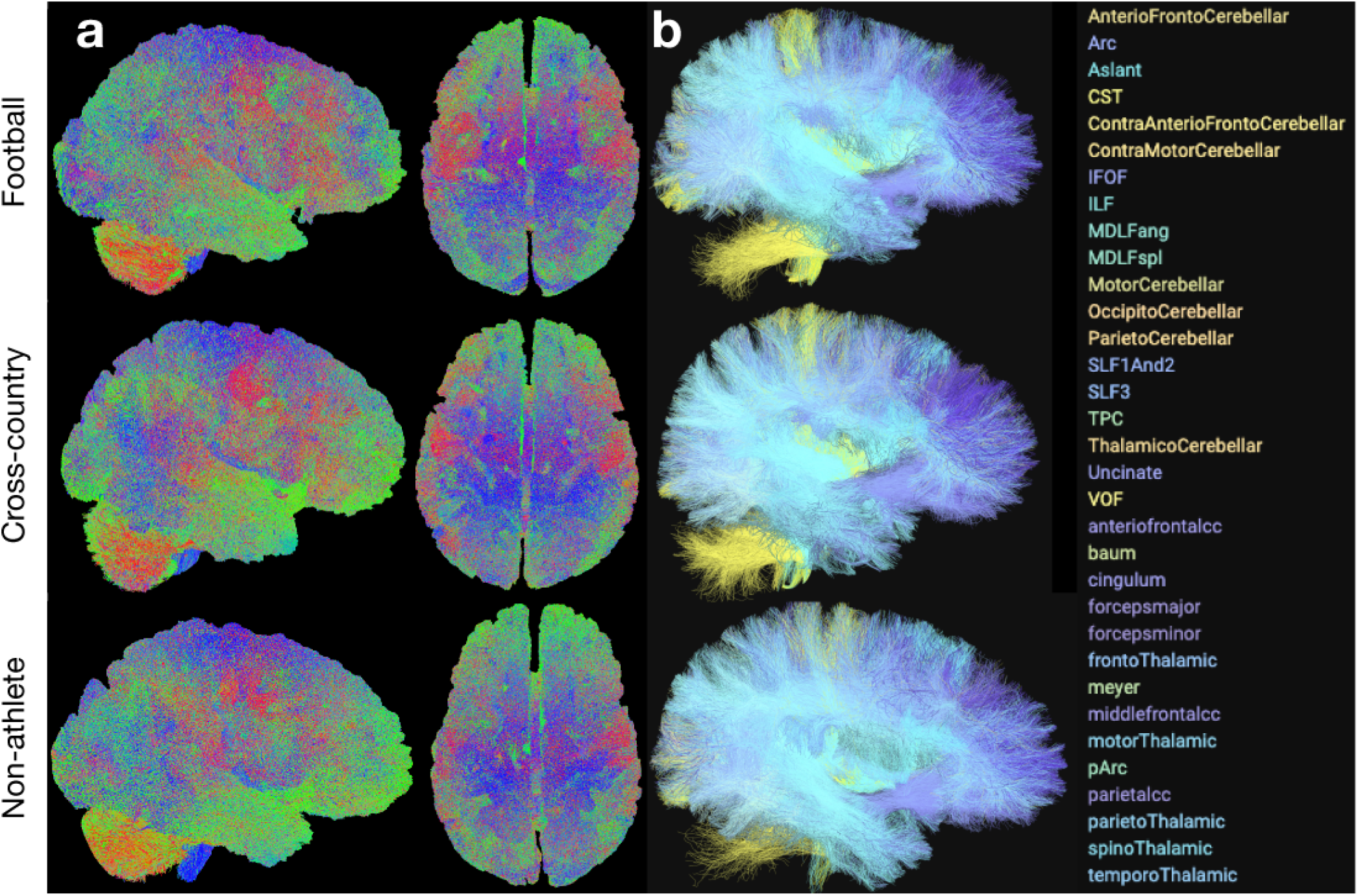
Ensemble Tractography. **a.** Example tractogram from each group subsampled from the ‘merged’ 3 million streamline tractogram (50k streamlines sampled per tractogram). The images were generated using https://doi.org/10.25663/brainlife.app.310. **b.** Example cleaned segmentation from each group. The images were generated using brainlife.io’s tract segmentation viewer.

### White matter microstructure modeling: Segmentation and cleaning

Following tractography, each participant’s whole-brain tractogram was segmented using a recently published methodology using anatomical definitions of common white matter tracts. In brief, this segmentation classifies streamlines as belonging to a particular tract based on their cortical terminations and known shape characteristics. See **Methods: White matter microstructure modeling (Segmentation & Cleaning)** for more details. Tract segmentation was performed using https://doi.org/10.25663/brainlife.app.188. Following segmentation, each tract was cleaned by removing outlier streamlines using https://doi.org/10.25663/brainlife.app.195. Each participant’s segmentation, both cleaned and uncleaned, is provided. **Figure 6b** provides representative white matter tract segmentation from participants from each group. From a qualitative perspective, each segmentation fills the whole-brain volume and contains a relatively high density of streamlines per tract. Each of the tracts is listed on the right. The images were generated using the brainlife.io tract segmentation viewer.

### White matter microstructure modeling: Tract profiles

Following white matter tract segmentation and cleaning, tract profilometry^87^ was performed for each participant and each tract. In brief, a central representation (i.e. ‘core’) of each tract was computed by weighted-average of the X,Y, and Z coordinates of all the streamlines in the tract. Streamlines were then resampled to 200 equally spaced nodes, and the average microstructural measures (DTI, NODDI) were computed at each node. The first and last ten nodes were removed, and then the profiles for each tract were averaged across each group. See **Methods: White matter microstructure modeling (Tract profiles)** for more details. Tract profiles were computed using https://doi.org/10.25663/brainlife.app.361. Tract profilometry data for all participants and tracts are provided. **Figure 7a** provides a representative Right ILF from a Football player. **Figure 7b** illustrates the centralized core of the ILF and the ODI values mapped along the tract. **Figure 7c** provides the group average ODI and FA tract profiles for the Right ILF. These profiles document the ability of this methodology for identifying group differences along a tract. Images were generated using the Matlab Brain Anatomy toolbox https://github.com/francopestilli/mba^86^ scripts available at https://github.com/bacaron/athlete-brain-study.

**Figure 7.**
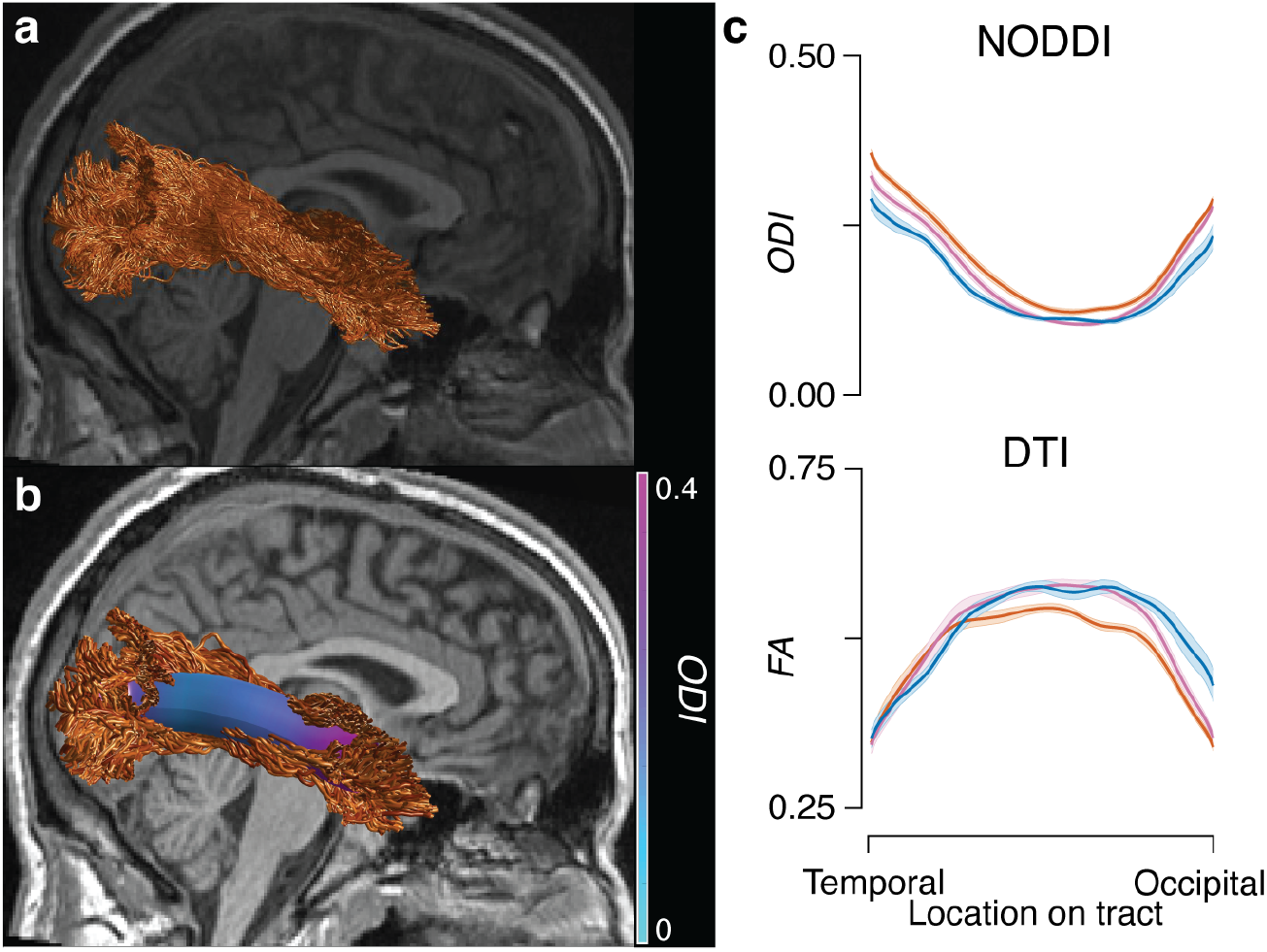
Tract Profiles. **a.** Example of a segmented Right ILF tract from a representative Football player. **b.** Example of the centralized ‘core’ representation of the Right ILF in the same subject as in **a**, with ODI mapped along the ‘core’. **c.** Group average tract profiles for ODI (*top*) and FA (*bottom*) for the Right ILF (orange: football players, pink: cross-country runners; blue: non-athletes; error bars ±1 SE.) Images were generated using the Matlab Brain Anatomy toolbox https://github.com/francopestilli/mba^86^ scripts available at https://github.com/bacaron/athlete-brain-study.

### White matter microstructure modeling: Network adjacency matrix generation

The whole-brain tractograms from each participant were used to generate structural connectomes. Specifically, measures of streamline count and density are computed between each node in the multimodal 180 cortical node parcellation and network matrices are generated ^94,95^. See **Methods: White matter microstructure modeling (Network generation)** for more details. Network adjacency matrices were generated using https://doi.org/10.25663/brainlife.app.394. **Figure 8** demonstrates group average connectivity matrices using log streamline count, log density, and average FA across the streamlines connecting nodes from each group and the total dataset. Images were generated using the *imagesc* function in MATLAB.

**Figure 8.**
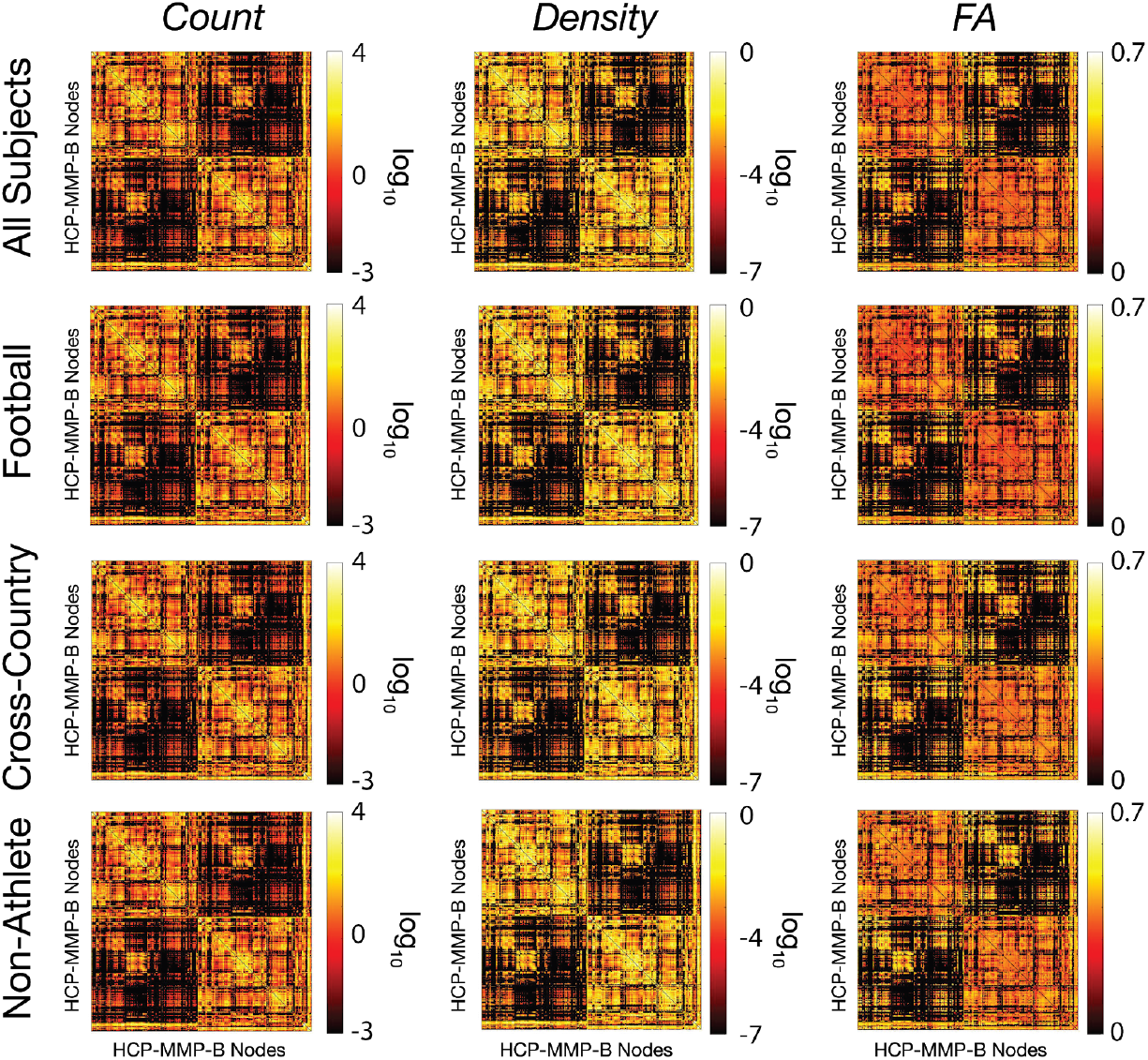
Average structural connectivity matrices. Twelve representative matrices of connectivity between brain regions defined in the 180 multimodal cortical atlas^66^ (i.e. HCP-MMP). Before averaging, any nodes in which half of the participants did not have a connection were removed. Adjacency matrices of average streamline count (*left*), density (*middle*), and FA (*right*) averaged across all subjects (*top*), football players (*2nd row*), cross-country runners (*3rd row*), and non-athlete students (*4th row*). Images were generated using *imagesc* in MATLAB.

### Cortical white matter microstructure mapping

Diffusion-based measures of microstructure were also mapped to a surface representation of the *midthickness* (i.e., the average coordinates between pial and white matter boundary surface; ^64^), here after simply referred to as ‘cortical.’ For more details on the differences between the two mappings, see **Methods: White matter microstructural modeling: DTI & NODDI**. The DTI and NODDI maps for each participant for the cortical white matter mapping analyses are provided as well. These measures were then mapped to the cortex following the procedures described in Fukutomi et al 2018^64^. See **Methods: Cortical white matter microstructure mapping** for more details. Diffusion measures were mapped using https://doi.org/10.25663/brainlife.app.379. Figure 9 demonstrates the quality of fit of DTI and NODDI measures on the cortical surface. Specifically, the FA and ODI maps mapped to a representative participant’s cortical surface. Anatomic landmarks, including higher FA and lower ODI in motor and somatosensory cortices, are consistent across participants and map well to the results presented in Fukutomi et al 2018. These images were generated using brainlife.io’s Connectome Workbench viewer.

**Figure 9.**
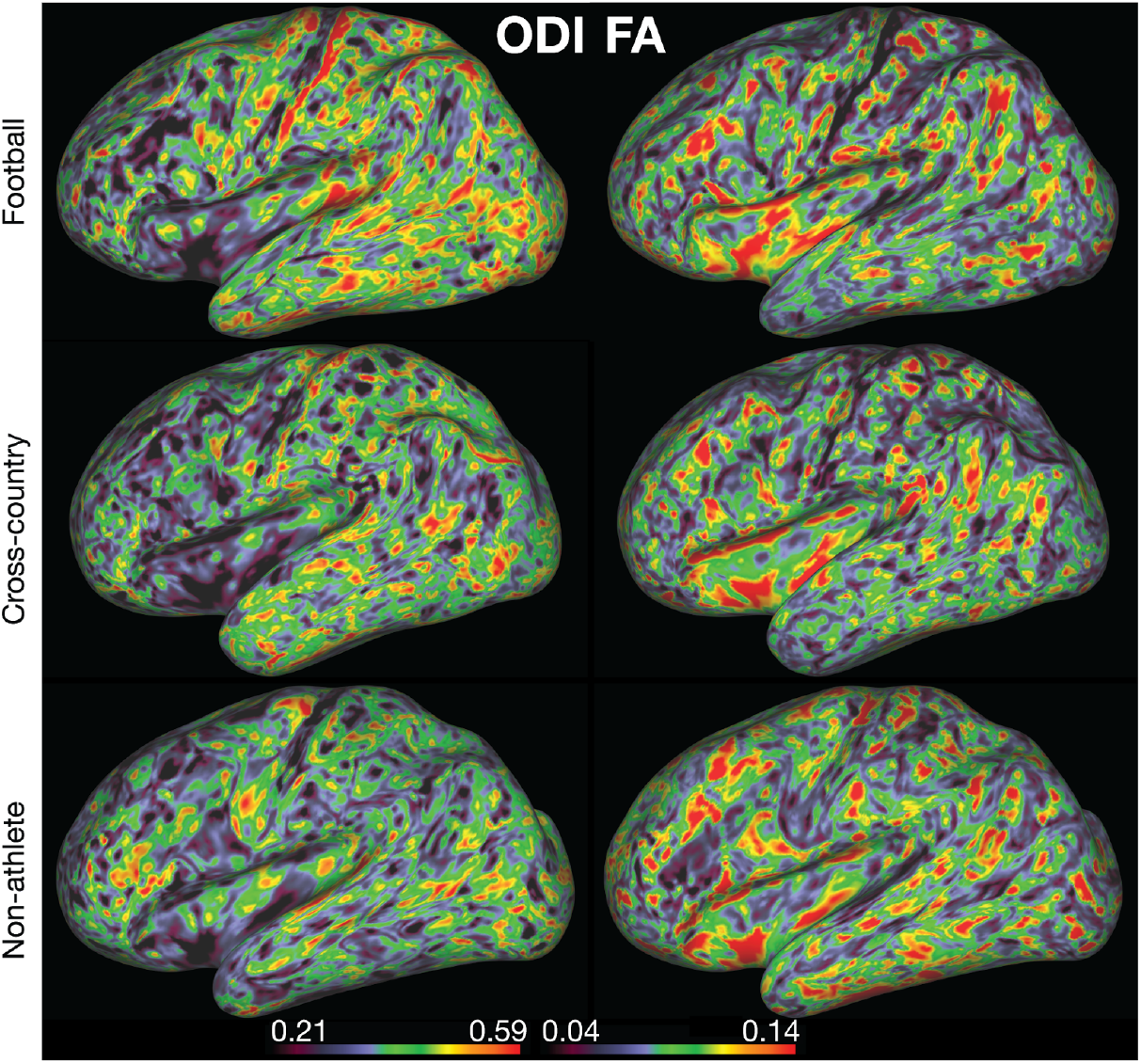
Surface-based mapping of microstructural white matter. Example ODI (left) and FA (right) estimates mapped to the cortical surface for a representative participant in each group (football: top, cross-country: middle, non-athlete: bottom). These images were generated using brainlife.io’s Connectome Workbench viewer.

### Mass and brain size

To further reduce the burden to full understanding of the dataset provided, we examined the potential differences between the groups in terms of mass and brain size. We collected data from each participant provided by the Freesurfer segmentations regarding total brain volume, cortical volume, white matter volume, and cortical thickness and computed one-way ANOVAs between our groups. We identified a significant difference in body mass between Football players and the other two groups (F(2,39), p < 0.0001; Figure 10e). However, we did not observe any significant effects of group on brain volume (Figure 10a), cortical volume (Figure 10b), white matter volume (Figure 10c), or cortical thickness (Figure 10d).

**Figure 10.**
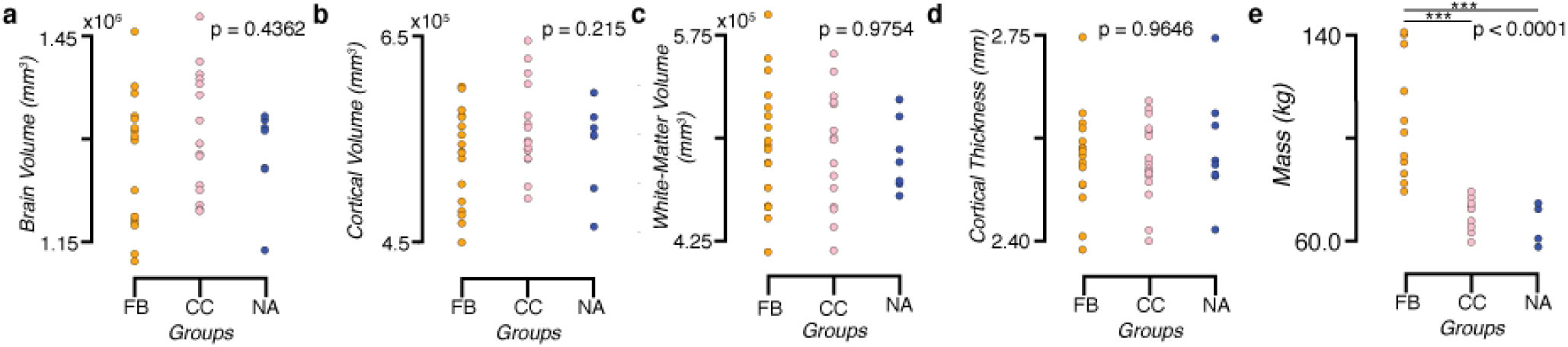
Total brain volume, gray -matter cortical volume, white matter volume, and average gray -matter cortical thickness show no differences between groups despite differences in body mass. Group distributions of total brain volume (**a**), cortical volume (**b**), white matter volume (**c**), and average cortical thickness (**d**). One-way ANOVAs showed no significant differences between the groups in these measures. A significant difference was observed between the mass (**e**) of football players and the two other groups (p < 0.005 Bonferroni corrected).

## Usage Notes

The data are publicly available on brainlife.io using the following DOI: https://doi.org/10.25663/brainlife.pub.14^93^. The following video shows how to access, download and visualize the data published with the record: https://www.youtube.com/watch?v=QEWEQydpbz4

Data files can also be downloaded, and some can be organized into BIDS standard ^92^. The data derivatives are stored in numerous formats, including NIFTI, TCK, GIFTI, and .mat. Access to the published data is currently supported via (i) web interface and (ii) Command Line Interface (CLI).

The brainlife.io CLI can be installed on most Unix/Linux systems using the following command:

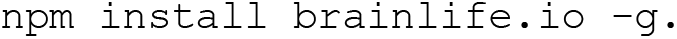

The CLI can be used to query and download partial or full datasets. The following example shows the CLI command to download all T1w datasets from a subject in the publication data:

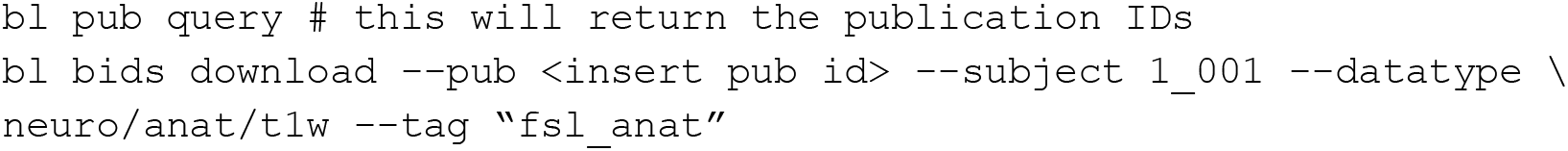

The following command downloads the data in the entire project (from Release 2) into BIDS format:

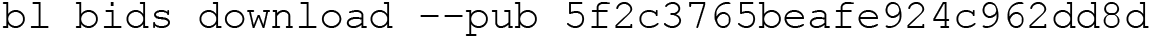

Additional information about the brainlife.io CLI commands can be found at https://github.com/brainlife/cli.

## Code availability

**Table 1** below reports the links to each web service and github.com URL implementing the processing pipeline. All code not found on brainlife.io, including visualization code, can be found at https://github.com/bacaron/athlete-brain-study.

## Acknowledgments

This research was supported by NSF OAC-1916518, NSF IIS-1912270, NSF IIS-1636893, NSF BCS-1734853, NIH NIDCD 5R21DC013974-02, NIH 1R01EB029272-01, NIH NIMH 5T32MH103213, NSF GRFP-1342962, the Indiana Spinal Cord and Brain Injury Research Fund, Microsoft Faculty Fellowship, the Indiana University Areas of Emergent Research Initiative “Learning: Brains, Machines, Children.” We thank Soichi Hayashi, and David Hunt for contributing to the development of brainlife.io, Craig Stewart, Robert Henschel, David Hancock and Jeremy Fischer for support with jetstream-cloud.org (NSF ACI-1445604). We also thank The Indiana University Lawrence D. Rink Center for Sports Medicine and Technology and Center for Elite Athlete Development for contributing funding to athletic scientific research and for the development of a new research facility.

## Author contributions

BC, RS, and FP wrote the manuscript text, performed analyses, and prepared figures 1–10. BC, DB, LK, JF, BM, and FP contributed to software development. BC, DK, HC, SN, NP, and FP contributed to data curation. All authors reviewed the manuscript.

